# Defining trait-based microbial strategies with consequences for soil carbon cycling under climate change

**DOI:** 10.1101/445866

**Authors:** Ashish A. Malik, Jennifer B. H. Martiny, Eoin L. Brodie, Adam C. Martiny, Kathleen K. Treseder, Steven D. Allison

**Author notes:** Correspondence: Ashish A. Malik.

## Abstract

Microorganisms are critical in terrestrial carbon cycling because their growth, activity and interactions with the environment largely control the fate of recent plant carbon inputs, as well as the stability of assimilated carbon (Gleixner, 2013; Schimel and Schaeffer, 2012). Soil carbon stocks reflect a balance between microbial decomposition and stabilization of organic carbon. The balance can shift under altered environmental conditions (Davidson and Janssens, 2006), and new research suggests that knowledge of microbial physiology may be critical for projecting changes in soil carbon and improving the prognosis of climate change feedbacks (Allison et al., 2010; Bradford et al., 2016; Liang et al., 2017; Wieder et al., 2015). Still, predicting the ecosystem implications of microbial processes remains a challenge. Here we argue that this challenge can be met by identifying microbial life history strategies based on an organism’s phenotypic characteristics, or traits, and representing these strategies in models simulating different environmental conditions.

What are the key microbial traits for microbial carbon cycling under environmental change? Microbial growth and survival in soil are impacted by multiple traits that determine responses to varying resource availability and fluctuating abiotic conditions (Wallenstein and Hall, 2012). Cellular maintenance activities include production of extracellular enzymes to degrade and acquire resources, biomolecular repair mechanisms, maintenance of cellular integrity, osmotic balance, and cell motility (Bradley et al., 2018; Geyer et al., 2016; Roller and Schmidt, 2015). It is conceivable that microbial investment into maintenance activities would be high in soils, with their highly heterogeneous and temporally variable resource distribution and stressful abiotic conditions like extremes of moisture, temperature, pH and salinity (Schimel et al., 2007).

Life history strategies represent sets of traits that tend to correlate due to physiological or evolutionary tradeoffs, with different strategies favoured under different environmental conditions. For example, metabolic investments in degradative enzyme production for resource acquisition can reduce the efficiency of cellular growth (Figure 1, Frank, 2010; Lipson, 2015). Furthermore, stress tolerance traits can trade off against investment in resource acquisition and growth yield (Figure 1, Schimel *et al.,* 2007; Manzoni *et al.,* 2014; Sinsabaugh *et al.,* 2013; Berlemont *et al.,* 2014). Although some stress tolerance mechanisms may have collateral benefits, the costs must generally be paid at the expense of other physiological processes if resources are limited. Ultimately, microbial metabolic investments and the resulting tradeoffs among growth yield, resource acquisition and stress tolerance determine the contribution of microbial processes to ecosystem level carbon fluxes. Thus, information about microbial strategies can be useful in linking microbial physiology to ecosystem function.

**Figure 1:**
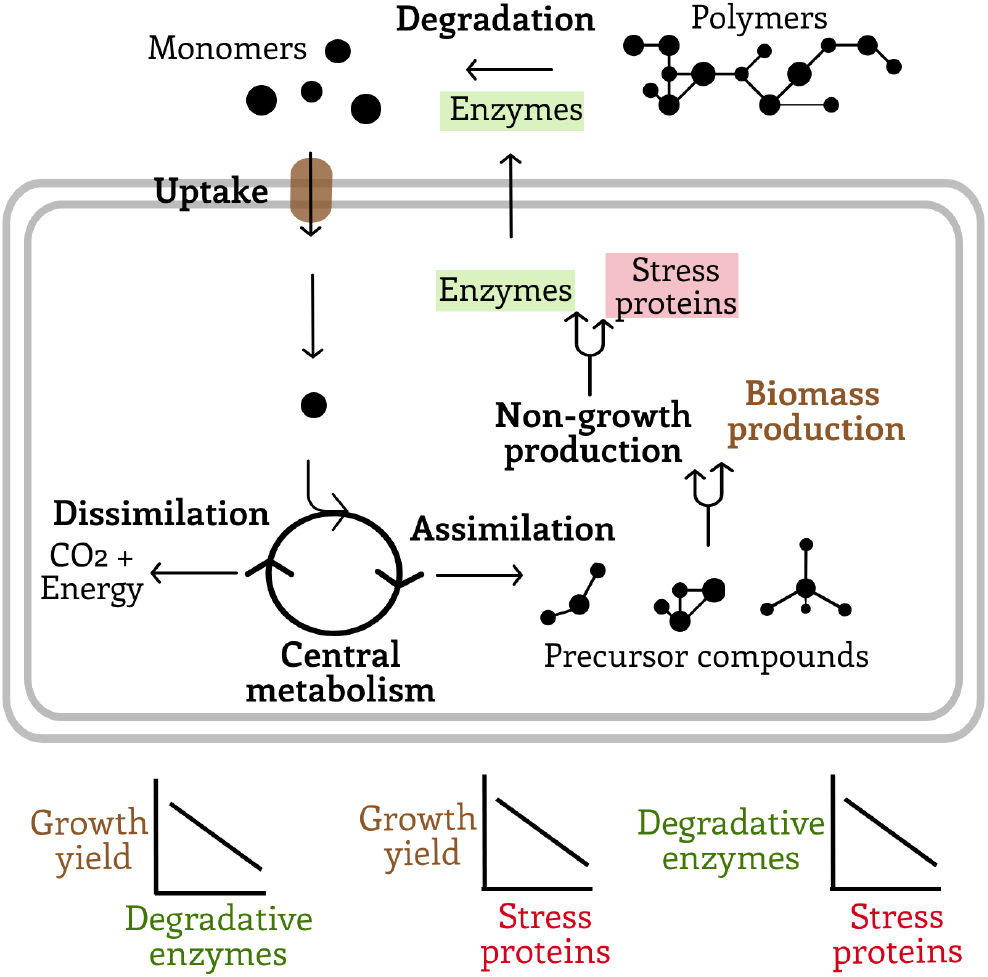
Schematic showing cellular C flux that includes depolymerisation, substrate uptake, assimilation, dissimilation, biomass synthesis and non-growth production. Extracellular enzyme production represents investment in resource acquisition, stress protein production is linked to stress tolerance mechanisms, and biomass production reflects higher growth yield. Forked arrows signify metabolic points where hypothesized tradeoffs in traits might occur. The expected empirical relationships among the key traits are also shown.

## Life history concepts in macroecology

In macroecology, trade-offs in key fitness traits have been represented through conceptual theories of r- and K-selection, the “leaf economics spectrum” and Grime’s competitor-stress tolerator-ruderal (C-S-R) framework. The r- and K-selection concept recognises two functional groups of organisms: r-selected strategies have short life expectancy and large reproductive effort whereas K-selected strategies have long life expectancy and invest a smaller proportion of energy and resources into reproduction (Pianka, 1970). Furthermore, leaf economics refers to the resource-driven tradeoffs among leaf traits that regulate photosynthesis (Reich et al., 1997; Wright et al., 2004). Grime’s C-S-R triangle is an alternative framework that enumerates three major plant life history strategies: competitors(C) excel at maximizing resource capture in productive and undisturbed systems, stress tolerators (S) prevail in continuously low-resource and stressful conditions, and ruderals (R) occupy recently disturbed but less stressful habitats (Grime, 1977). Such tradeoffs have been shown to apply globally across biomes thus providing the quantitative basis to functionally represent the enormous taxonomic diversity of plant communities in vegetation models (Reich et al., 1997; Wright et al., 2004).

## Applying trait-based theories of life history in microbial ecology

Following on macroecological theory, microbial ecologists have proposed trait-based classifications of microorganisms. The copiotroph-oligotroph continuum was proposed as analogous to the r- and K-selection theory for plants and animals (Fierer et al., 2007; Koch, 2001). This classification was mostly based on microbial substrate preferences and growth rates and has since been widely applied in various contexts (Fierer et al., 2012; Thomson et al., 2013). Several recent efforts have also applied C-S-R life history strategies to microbial systems, particularly in the context of anthropogenic environmental change (Fierer, 2017; Ho et al., 2013; Krause et al., 2014; Wood et al., 2018). Ho *et al.* (2013) classified methane-oxidising bacteria into C-S-R life strategies based on activity, recovery from disturbances, substrate utilization patterns and stress tolerance. Krause *et al.* (2014) later generalized the same framework for all bacteria while emphasizing that additional experiments would be needed to verify the microbial strategies and their underlying traits. Wood *et al.* (2018) justified a microbial C-S-R classification based on different traits derived from predicted functional datasets aimed at assessing the impact of cadmium and influence of the rhizosphere on microbial community assembly. These previous studies justify additional theory development and experimental evidence to validate the C-S-R framework in microbial ecology.

Although a general theory of life history is attractive, the C-S-R strategies do not necessarily map clearly on to microbial systems. In the plant C-S-R framework, Grime (1977) defined habitats based on gradients in disturbance intensity and stress, including multiple abiotic and resource-based factors. These gradients were thought to select for C, S, or R strategies defined by plant traits including morphology, growth form, relative growth rate, leaf longevity, phenology, and seed production. Although some microbial traits like growth rate, biomass turnover, and dormancy may be analogous to plant traits, it can be challenging to apply a theory about autotrophic macroorganisms to heterotrophic microbes. Thus, it remains unclear how C-S-R strategies emerge from underlying microbial traits. It is notable, however, that Grime (1977) himself suggested some traits useful in mapping fungi to the C-SR strategies: rapid growth and soluble carbohydrate use for ruderals, dense mycelium and rhizomorph production for competitors, and slow growth, persistent mycelium, and low spore production for stress tolerators.

Moreover, given the vast metabolic diversity of microorganisms and the complexity of functional omics datasets, it is also unclear what dimensionality is needed to adequately describe microbial life history strategies (Laughlin, 2013). A three-dimensional trait space much like the C-S-R framework is a good start in advancing trait-based microbial ecology, while keeping in mind that increasing trait dimensionality may help better predict species distribution in a trait space.

Here we propose a revised life history theory for microbes that builds on the work by Wood *et al.* (2018). Their framework justifies microbial C-S-R classifications based on predicted genomic traits. Traits of the competitor strategy focus on antibiotic production and resource acquisition through siderophores and membrane transporters. Stress-tolerator traits relate to damage repair and maintenance of cell integrity. Ruderal microbial traits include investment in processes that promote rapid growth. Wood et al. also define a fourth group of traits related to foraging, such as chemotaxis and flagellum production. Our revised framework emphasizes three strategies somewhat analogous to Wood et al.’s version of C-S-R but reclassified to emphasize the link between carbon cycling and microbial physiology. We propose three main microbial life history strategies: high yield (Y), resource acquisition (A) and stress tolerance (S), or Y-AS (Figure 2A).

**Figure 2.**
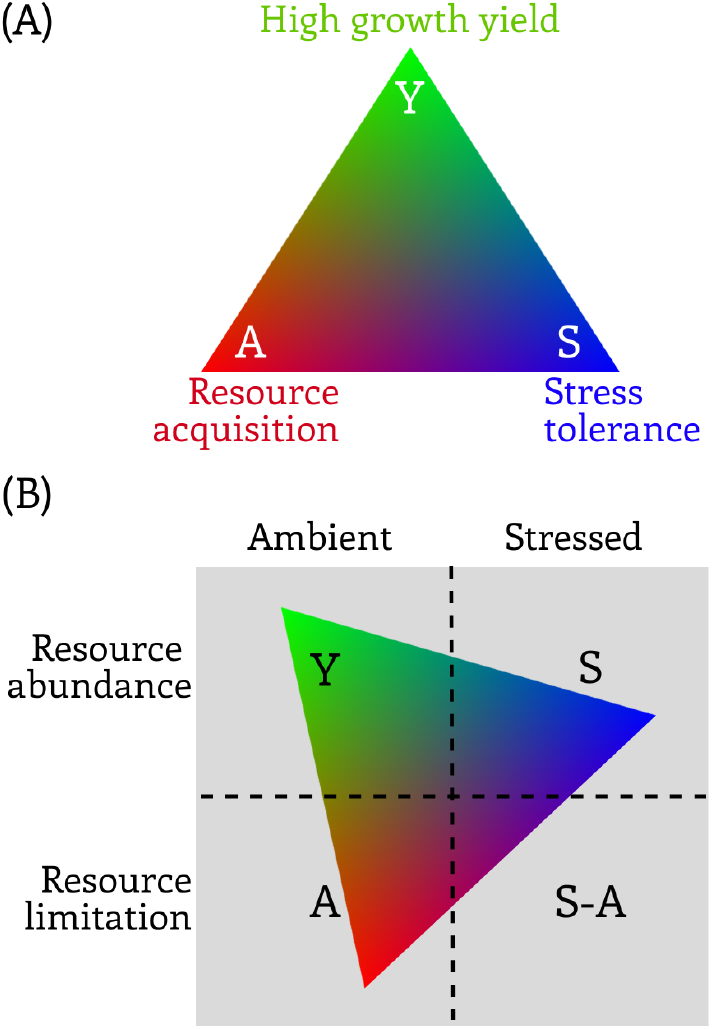
A) Conceptual figure of microbial Y-A-S life history strategies. High yield (Y): maximises growth efficiency as a result of reduced investments in stress tolerance and resource acquisition; resource acquisition (A): preferential investment in cellular resource acquisition machinery; stress tolerance (S): preferential investment in stress tolerance mechanisms. B) Hypothesized strategies favoured under particular treatment combinations. The microbial three-dimensional Y-A-S triangle is arrayed on the combinations.

## The High Yield (Y) strategy

We define yield as the amount of microbial biomass produced per unit of resource consumed (Hagerty et al., 2014; Manzoni et al., 2012). High yield strategists maximize the fraction of resource uptake that is allocated to growth processes by investing in central metabolism and associated assimilatory pathways such as amino acid, nucleotide, and fatty acid synthesis. Conditions of high resource availability and minimum stress are expected to favour the high yield strategy (Figure 2B). Although parallel to the plant ruderal strategy, the Y strategy is not defined by growth rate, i.e. the change in microbial biomass per unit time (Bradley et al., 2018; Geyer et al., 2016; Roller and Schmidt, 2015). *In-situ* growth rate is not a coherent strategy but rather a complex emergent property that depends on both growth yield and the rate of resource acquisition. In fact, evidence suggests that growth rate and yield may have a negative relationship (Muscarella and Lennon, 2018; Roller et al., 2016) or a positive one (Ng, 1969) depending on the system.

Growth rate and yield may also diverge within the Y strategy such that high yields are achieved with different growth rates. Although *in-situ* growth rate is an emergent property, the maximum potential growth rate (μ_max_) is a physiological trait that varies across microbial taxa (Dolan et al., 2017). There is some evidence for a rapid growth, low yield strategy characterised by enhanced metabolism, large cell sizes, high ribosomal production, and high maximum growth rates (Roller et al., 2016). However, the strength of the growth rate-yield tradeoff is somewhat inconsistent across individual, population, and community levels (Malik et al., 2018; Muscarella and Lennon, 2018; Roller et al., 2016; Van Bodegom, 2007). Overall, we argue that high yield is a coherent, trait-based strategy whereas high growth results from combining any strategy with the right environmental conditions.

## The Resource Acquisition (A) strategy

Our resource acquisition strategy replaces the plant competitor strategy because microbial competition is mainly over resources. In fact, one could also argue this is true for plants. In soils, microorganisms produce extracellular enzymes to break down complex resources (Table 1, Allison et al., 2010; Frank, 2010; Lipson, 2015). Thus, resource acquisition by heterotrophic microbes depends on uptake of depolymerised substrates using various membrane transporters (Table 1). The level of investment in extracellular enzyme production often reflects substrate status (quality and quantity) of the local environment (Allison and Vitousek, 2005). Wood et al.’s foraging traits can readily be assimilated into our resource acquisition strategy. However, contrary to Wood et al.’s hypothesis that investment in resource acquisition traits is higher in resource abundant environments like the rhizosphere, we propose that this strategy will prevail in low resource conditions where microbes would be under selection to increase resource capture at the expense of growth yield. It is also likely that organisms have acquisition strategies that are either uptake optimised (when precursors compounds are readily available in the environment) or depolymerization optimised (when resources are scarce and complex) or a combination of both (Malik et al., 2018; Zhalnina et al., 2018).

**Table 1.**
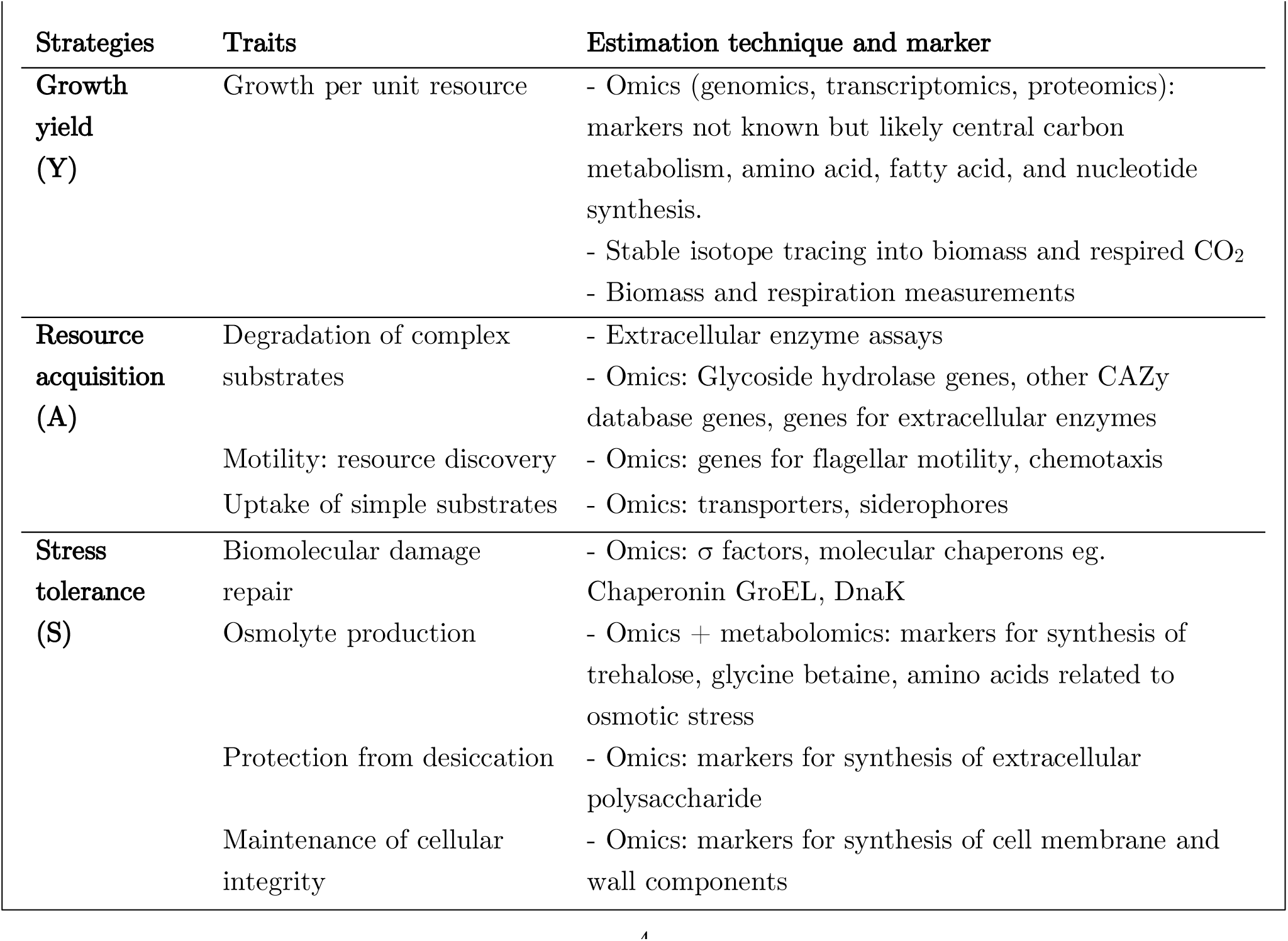
Y-A-S strategies, underlying traits and tools to extract trait information.

## The Stress Tolerator (S) strategy

We adopt Grime’s stress tolerance strategy with little modification because it aligns well across plants and microbes. Soil microbes experience a variety of stressors that change their physicochemical environment and to which they respond through physiological acclimation mechanisms in order to survive and grow (Schimel et al., 2007). Specific traits for stress tolerance depend on the kind of abiotic stress experienced by microbial communities. Regardless of the form of stress imposed, certain global patterns in phenotypic expression are common, including σ factors or molecular chaperons aimed to minimise or mitigate biomolecular damage (Table 1, Finn et al., 2016; Hecker and Völker, 2001; Malik et al., 2017; Wood et al., 2018). In cases such as high acidity or salinity, microbes employ various strategies to maintain cellular integrity and osmotic balance through changes in the structure and composition of cell envelopes (Wood, 2015). Under drought scenarios, stress tolerance strategies involve production of osmolytes like trehalose and glycine betaine or synthesis of extracellular polymeric substances (EPS)— usually polysaccharides—to protect cells from desiccation (Table 1, Bouskill et al., 2016; Schimel et al., 2007). Thus, microbes exposed to suboptimal abiotic conditions would possess traits linked to stress tolerance at the expense of other traits.

## Strategies under varying conditions

Tradeoffs in resource allocation among the Y-A-S strategies should prevent microbes from excelling at multiple strategies. Different strategies should be favoured under different environmental conditions arising from spatial or temporal variability in resource status and abiotic conditions (Figure 2B). For example, resource limitation or abiotic stress should select against Y-strategies because of a need for investment in costly resource acquisition or stress-tolerance mechanisms. In environments with high availability of polymeric resources (e.g. cellulose) but few simple resources (e.g. glucose), A-strategists should outcompete Y-strategists by investing in extracellular enzyme machinery. Thus, A-strategists catalyse polymer decomposition and soil carbon loss, whereas Y-strategists may convert monomeric substrates into more stable microbial biomass residues (Kallenbach et al., 2016). These examples illustrate how life history tradeoffs can have consequences for soil carbon dynamics.

## Approaches for measuring and testing Y-A-S strategies

Technological innovations like next generation sequencing have massively improved our understanding of the taxonomic and functional diversity of soil microbial communities and their shifts in response to anthropogenic influences (Fierer, 2017). Current approaches have mostly focused on identifying taxonomic and functional responses to environmental changes. However, integration of these large microbial molecular datasets with process rate measurements remains a challenge, thereby making it difficult to link microbial diversity and function with ecosystem processes (Krause et al., 2014; Rocca et al., 2015). More efforts are needed to determine how microbial genomic information translates into traits that influence abundance, metabolite production, and ultimately carbon cycling rates in ecosystems.

A variety of omic tools are now available to assess the functional response of microbes to environmental change. However, information overload presents challenges for incorporation of trait data into ecosystem models. This challenge can be met by simplifying the data through the Y-A-S framework and fitting genes, transcripts, proteins and metabolites into our hypothesised three-dimensional trait space (Fierer, 2017; Krause et al., 2014). Population-level trait information can be gathered from sequenced genomes where cultured microbial strains are available (Zhalnina et al., 2018). In other cases, individual or population genomes can be assembled from culture-independent shotgun metagenomic datasets; this novel approach is gaining popularity as it facilitates physiological investigations of hitherto uncultured taxa (Hu et al., 2016). In addition, traits can be measured at the community level to integrate tradeoffs across populations and characterise trait impacts on ecosystem function (Geyer et al., 2016; Hall et al., 2018). Many phenotypic traits can also be measured directly, even if the underlying genetic mechanisms are complex and cannot be determined.

## Omics and physiological techniques to quantify traits

Growth yield (synonymous with growth efficiency) is a challenging property to extract from omics datasets because we still do not understand its genetic determinants. However, there are quantitative methods for estimating growth yield and its components (Table 1, Geyer *et al.,* 2016). Approaches include measuring the change in biomass proxies and respiratory loss or following a tracer—commonly a stable isotope— in cellular fractions. Yield is often measured as the proportion of C substrate invested into biomass relative to that lost through respiration.

Recent studies emphasize, though, that growth yield is actually an emergent and dynamic property of multiple underlying traits related to cellular maintenance, protein synthesis and export, cellular stoichiometry, and respiratory pathways (Hagerty et al., 2018; Manzoni et al., 2012). This complexity creates challenges for measuring growth yield but emphasizes the trait tradeoffs inherent in the high-yield (Y) strategy. Increased respiration associated with enzyme production (A strategy) or maintenance of cellular integrity (S strategy) should directly and negatively affect measured growth yield.

Resource acquisition traits have been estimated with omics and biochemical techniques at both the population and community levels (Table 1). Extracellular enzyme assays provide estimates of microbial enzyme activity and the potential to degrade various complex substrates. Genes and transcripts encoding these enzymes can also be predicted from omics datasets. In addition, there is growing interest in linking Carbohydrate-Active Enzyme (CAZy) database genes to microbial substrate degradation and resource acquisition potential. The CAZy database includes genes that code for enzymes that synthesise and break down complex carbohydrates and glycoconjugates (Cantarel et al., 2009). For example, glycoside hydrolases (GH) are involved in plant cell wall degradation and act on glycosidic bonds between carbohydrates or between carbohydrates and non-carbohydrate moieties (Berlemont, 2017; Naumoff, 2011). Once the complex polymers are degraded into simpler molecules, they are taken up by transporters (Figure 1). A variety of transporters, particularly the ATP-binding cassette transporters (ABC-transporters), with differential substrate specificity can also be predicted from omics datasets. Greater investment in uptake transporters has been observed in root-associated microorganisms with plentiful substrates allowing the cells to reduce investment into extracellular enzyme production and increase their growth yield (Malik et al., 2017; Zhalnina et al., 2018).

Stress tolerance traits in the form of σ factors, molecular chaperons or specific physiological adaptations can be extracted from widely used omics tools (Finn et al., 2016; Malik et al., 2018). Some low molecular weight metabolites synthesised in response to environmental stimuli can be quantified using mass spectrometry tools like LC-MS and FT-ICRMS (Table 1, Tfaily *et al.,* 2015; Swenson *et al.,* 2018; Bouskill *et al.,* 2016).

## Carbon cycling implications of tradeoffs in Y-A-S traits

We posit that microbial physiological investments and the resulting tradeoffs among key traits determine the contribution of microbial processes to ecosystem level carbon fluxes. Microbial physiological responses and the resulting effects on growth yield can affect carbon balance through two main mechanisms. On the one hand, microbial biomass is thought to contribute significantly to organic matter accumulation and hence to the genesis of soil organic matter (Gleixner, 2013; Kallenbach et al., 2016). On the other hand, microbial biomass and extracellular enzymes contribute to plant litter and soil organic matter degradation.

Under our Y-A-S framework, Y-strategists with increased investment into growth and biomass production would contribute to microbial residue formation that can be stabilized in the soil organo-mineral matrix. In contrast, A-strategies should contribute more to decomposition and carbon loss through investment in extracellular enzyme production (Kallenbach et al., 2016; Schimel and Schaeffer, 2012). Carbon impacts of S-strategists might depend on the type of stress compounds produced, with more complex compounds like extracellular polymeric substances (EPS) contributing more to carbon storage than simple compounds like osmolytes (Bouskill et al., 2016; Schimel et al., 2007). By diverting investments away from growth, S-strategists could also reduce soil carbon accumulation. The effect of microbial physiological adaptation to climate change and its consequences for soil C cycling could thus be determined by assessing shifts in microbial Y-A-S life history strategies.

## Approaches to modelling Y-A-S strategies to predict carbon fluxes

Representing microbial diversity has been a big challenge for models projecting ecosystem responses to environmental change (Hall et al., 2018). This challenge introduces uncertainty that affects model predictions of future climatic change (Bradford et al., 2016; Sulman et al., 2018). Such uncertainties imply a need for better mechanistic models, and improved representation of microbial diversity and physiology could increase the accuracy of projected soil carbon fluxes.

Previously, functional traits have been incorporated into ecosystem models like the MIcrobial-MIneral Carbon Stabilization (MIMICS) model to predict the biogeochemical response of soil organic matter decomposition and stabilization (Wieder et al., 2015). In this model, copiotrophic and oligotrophic functional groups represent fast-growing low yield and slow-growing high yield strategists, respectively. However, this classification may be insufficient to accommodate the vast metabolic flexibility of soil microbial populations (Fierer, 2017; Krause et al., 2014). In addition, traits for acquisition of complex resources and tolerance to abiotic stressors are difficult to incorporate into the copiotrophic-oligotrophic dichotomy (Schimel et al., 2007; Schimel and Schaeffer, 2012).

Benefiting from the accumulation of trait data (Reich et al., 1997; Sakschewski et al., 2015; Wright et al., 2004), trait-based approaches are emerging in vegetation modelling as a paradigm shift from approaches based on aggregated plant functional types (Scheiter et al., 2013). For example, a trait-based vegetation model was successfully applied to uncover the roles of trait diversity in conferring resilience to the Amazon forests under climate change (Sakschewski et al., 2016, 2015). The model implemented tradeoffs among five major plant traits that determine growth and mortality of individual trees.

Likewise, a trait-based modelling framework based on microbial Y-A-S strategies holds promise for representing the immense diversity of microbial communities in simulations of system-level processes at various spatial scales (Hall et al., 2018). The cellular mechanisms underlying tradeoffs in key physiological traits can be incorporated into microbial functional models like MIMICS to reveal how these tradeoffs structure microbial communities and their resulting carbon cycle functions. Thus, embracing microbial complexity through the trait-based modelling approach, facilitated by the omics-derived data, would enable better predictions of ecosystem processes at larger temporal and spatial scales.

Models representing continuous variation in traits across taxa are particularly promising tools for predicting biogeochemical processes based on the Y-A-S framework. For example, DEMENT is a local scale, trait-based model that simulates litter decomposition and soil carbon transformations by diverse microbial communities (Allison, 2014, 2012). The model uses relationships between Y and A traits as a mechanistic basis for predicting how microbial communities and carbon cycling processes will respond to future environmental change (Figure 3, Webb *et al.,* 2010). Yield in the model is a function of multiple factors, including substrate type and stoichiometry, enzyme production rates, uptake investment, and temperature. The most recent version of DEMENT also includes a simple representation of drought stress tolerance (Allison and Goulden, 2017). After incorporating trait tradeoffs derived from omics or other data sources for individual taxa, DEMENT projects community responses and carbon cycling consequences under simulated environmental conditions. Model outputs can be validated with *in-situ* trait distributions at a community level or with ecosystem processes like organic matter decomposition rates (Figure 3). This validation approach can also be applied to other individual-based models that simulate spatial structuring of microbial populations based on functional groups characterised by traits (Ginovart et al., 2005; Kaiser et al., 2014). Based on the successful trait-based modelling of global vegetation, we expect rapid progress in developing local- to global-scale models that incorporate microbial traits.

**Figure 3.**
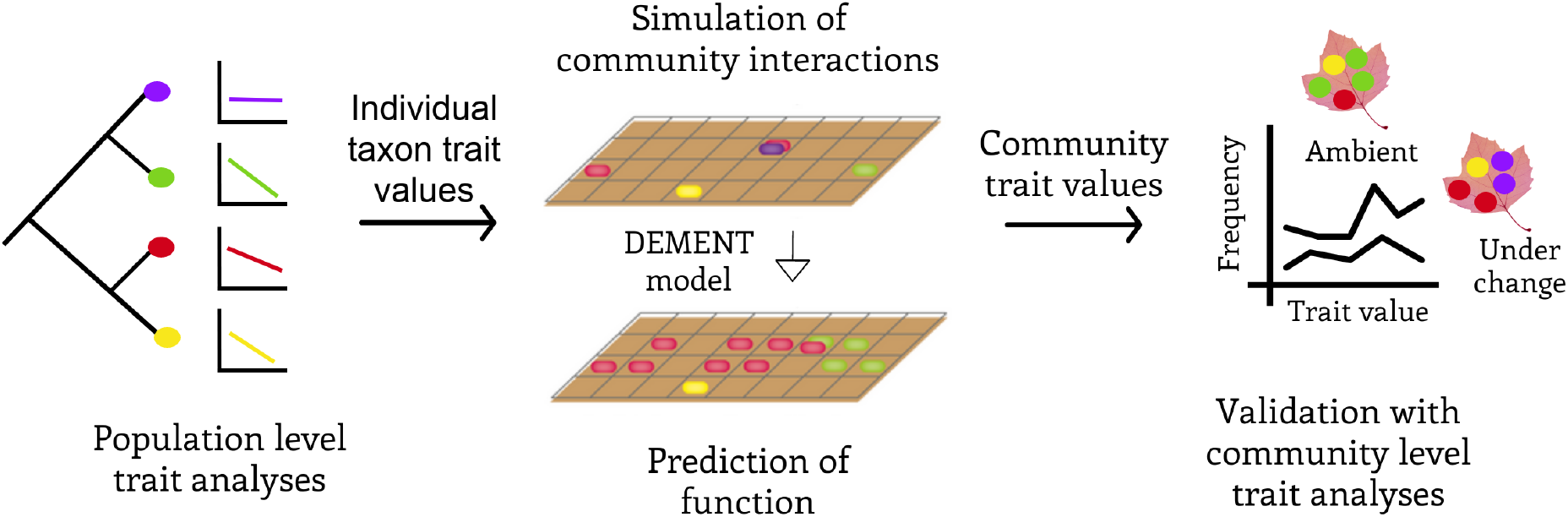
Summary of the proposed trait-based framework incorporating microbial life history strategies into the DEMENT model to predict community response and its ecosystem consequences under environmental change (adapted from Allison and Goulden, 2017).

## Conclusions

There is growing interest in applying trait-based concepts to predict the microbial mechanisms driving global biogeochemical cycles. By adapting several theories from macroecology, we define microbial high yield, resource acquisition, and stress tolerator strategies based on key traits that are linked to organismal fitness. Our Y-A-S framework can guide new empirical and modelling studies on the mechanisms driving soil carbon fluxes. We anticipate that these approaches will improve our understanding of the physiological constraints facing microbes under anthropogenic influence. By linking population-level response traits to community and ecosystem processes, our life history theory can improve predictive understanding of soil C responses to future climatic change.

## Acknowledgements

We acknowledge funding from the US DOE Genomic Science Program, BER, Office of Science project DE-SC0016410.

